# Risk of bias in Cochrane systematic reviews: assessments of risk related to attrition bias are highly inconsistent

**DOI:** 10.1101/366658

**Authors:** Andrija Babic, Ruzica Tokalic, João Amílcar Silva Cunha, Ivana Novak, Jelena Suto, Marin Vidak, Ivana Miosic, Ivana Vuka, Tina Poklepovic Pericic, Livia Puljak

**Author notes:** Corresponding author: Prof. Livia Puljak, MD, PhD.

## Abstract

**Background:** An important part of the systematic review methodology is appraisal of the risk of bias in included studies. Cochrane systematic reviews (CSRs) are considered golden standard regarding systematic review methodology, but Cochrane’s instructions for assessing risk of attrition bias are vague, which may lead to inconsistencies in authors’ assessments. The aim of this study was to analyze consistency of judgments and support for judgments of attrition bias in CSRs of interventions published in the Cochrane Database of Systematic Reviews (CDSR).

**Methods:** We analyzed CSRs published from July 2015 to June 2016 in the CDSR. We extracted data on number of included trials, judgment of attrition risk of bias for each included trial (low, unclear or high) and accompanying support for the judgment (supporting explanation). We also assessed how many CSRs had different judgments for the same supporting explanations.

**Results:** In the main analysis we included 10292 judgments and supporting explanations for attrition bias from 729 CSRs. We categorized supporting explanations for those judgments into four categories and we found that most of the supporting explanations were unclear. Numerical indicators for percent of attrition, as well as statistics related to attrition were judged very differently. One third of CSR authors had more than one category of supporting explanation; some had up to four different categories. Inconsistencies were found even with the number of judgments, names of risk of bias domains and different judgments for the same supporting explanations in the same CSR.

**Conclusion:** We found very high inconsistency in methods of appraising risk of attrition bias in recent Cochrane reviews. Systematic review authors need clear guidance about different categories they should assess and judgments for those explanations. Clear instructions about appraising risk of attrition bias will improve reliability of the Cochrane’s risk of bias tool, help authors in making decisions about risk of bias and help in making reliable decisions in healthcare.

## Introduction

Cochrane systematic reviews (CSRs) are produced using rigorous and evolving methodological standards and are therefore considered the gold standard when it comes to synthesis of evidence. The Cochrane has been at the forefront of applying the methods of evidence-based medicine (EBM) in the treatment and management of various conditions [1].

An important part of the systematic review methodology is appraisal of the risk of bias (RoB) in included studies. The potential effect of bias is that trialists will reach wrong conclusions about efficacy and safety of studied interventions. Bias can, therefore, negatively affect the estimated intervention effects [2].

In Cochrane systematic reviews RoB is appraised using Cochrane RoB tool, which has seven domains. Random sequence generation is analyzed as a potential selection bias, assessing potentially biased allocation to interventions due to inadequate generation of a randomised sequence. Allocation concealment can lead to another selection bias, potentially leading to biased allocation to interventions due to inadequate concealment of allocations prior to assignment. Blinding of participants and personnel is associated with performance bias due to knowledge of the allocated interventions by participants and personnel during the study. Blinding of outcome assessment, if done inadequately, can lead to detection bias due to knowledge of the allocated interventions by outcome assessors. Incomplete outcome data can yield attrition bias due to amount, nature or handling of incomplete outcome data. Selective reporting can cause reporting bias due to selective outcome reporting. And finally, there is a domain of RoB assessment called „other bias“, which is bias due to problems not covered elsewhere in the first six domains [3].

In the Cochrane RoB tool, the authors need to provide judgment about whether this risk is high, unclear or low for each domain. Furthermore, each judgment needs to be accompanied with a supporting explanation called ‘support for judgment’, which “describes what was reported to have happened in the study, in sufficient detail to support a judgement about the risk of bias.” The aim of the support for judgment is to ensure transparency about how these judgments about the level of risk of bias were reached [3].

The Cochrane Handbook provides vague instructions about assessing attrition bias, which may lead to inconsistent use of supporting explanations for judgments of attrition bias that one can find in CSRs. The aim of this study was to analyze whether Cochrane authors use consistent judgments for different supporting explanations of attrition bias in intervention CSRs published in the Cochrane Database of Systematic Reviews (CDSR).

## Methods

### Study design

Cross-sectional meta-epidemiological study of published CSRs was conducted.

### Inclusion and exclusion criteria

Intervention CSRs published from July 2015 to June 2016 were included by using Advanced search in The Cochrane Library. We excluded diagnostic CSRs, empty CSRs, overviews of systematic reviews and CSRs withdrawn in this period and CSRs that included only non-randomized studies. If the CSRs included randomized, quasi-randomized and non-randomized studies, we analyzed attrition bias in the RoB tables for the randomized studies only. CSRs that had multiple attrition bias judgments assessed for different outcomes in the same study were rare; therefore we reported them separately in order to better describe that methodological approach.

### Screening

Two authors independently assessed all titles/abstracts to establish eligibility of CSRs for inclusion. Discrepancies in judgment were resolved by the third author (LP).

### Data extraction

Data extraction table was developed and piloted using five CSRs. One author extracted data independently and the other author (AB) checked 10% of the extractions randomly. Discrepancies in data extraction were resolved by the third author (LP).

The following data were extracted: number of included trials, judgment of attrition risk of bias for each included trial (low, unclear or high) and accompanying ‘support for judgment’. To avoid terminological confusions, instead of ‘support for judgment’ hereby se use the expression ‘supporting explanation’. We also assessed how many CSRs had inconsistent judgments for the same supporting explanations (i.e. whether they had different judgments for the same supporting explanations). In the main analysis we reported only analysis of attrition bias for included CSRs with a single judgment, regardless of the number of supporting explanations that were provided for that judgment.

In the secondary analysis we investigated i) characteristics of attrition bias reporting for CSRs that reported multiple judgments of attrition bias for the same trial, ii) characteristics of risk of bias reporting in CSRs that did not have attrition bias domain, and iii) characteristics of risk of bias judgment reporting in CSRs that did not provide judgment in the form of “low, unclear and high”. Specific CSRs are marked in the body of this manuscript with the serial number of the downloaded record (for example, CSR #1).

### Statistics

Descriptive statistics was performed and data presented as frequencies and percentages. Data were analyzed using Microsoft Excel (Microsoft, Inc., Redmond, WA, USA).

## Results

Among 955 Cochrane systematic reviews published from July 2015 to June 2016 we included 729 CSRs in the main analysis. In the 729 included CSRs there were 1-105 included studies (median: 8 studies). In those CSRs we found 10292 attrition bias domains with single judgment about whether the CSR authors found this bias to be low, unclear or high. Although there was a single judgment, 3504/10292 (34%) supporting explanations contained more than one type of explanations related to risk of attrition bias. We categorized these different types of supporting explanations into four categories: percent of attrition in the RCT groups with higher attrition, difference in attrition between the groups, reporting of reasons for attrition and statistical comments. Only 27/10292 (0.26%) of supporting explanations had all four categories of explanations.

### First category: percent of attrition in the RCT groups with higher attrition

In the first category a third of supporting explanations were unclear (32%). The next most common type of supporting explanations were mentioning only total attrition (16%), indicating there was no attrition (15%) in the trial, providing only number of patients without a percent (11%), or indicating that attrition was not reported in a trial (8.8%) (Table 1).

**Table 1.** Number of explanations in a category for percent of attrition per group

| **Category for percent of attrition per group** | **N (%)** |
| --- | --- |
| Unclear | 3272 (31.8) |
| Total attrition only mentioned; attrition per group not reported | 1593 (15.5) |
| No attrition | 1544 (15) |
| Only number of patients, no percent provided | 1115 (10.8) |
| Not reported | 901 (8.8) |
| No explanation for this category | 414 (4) |
| 10-20% | 359 (3.5) |
| Above 30% | 276 (2.7) |
| Under 10% | 267 (2.6) |
| 20-30% | 216 (2.1) |
| Total attrition reported as percent; attrition per group reported as absolute numbers so it was not possible to judge percent attrition per group | 248 (2.4) |
| Information about attrition provided for one group only | 35 (0.3) |
| ‘Support for judgment’ box was blank: no explanation provided for the judgment | 27 (0.3) |
| Above certain percentage that is not precisely defined | 13 (0.1) |
| Under certain percentage that is not 10% | 6 (0.06) |
| There was no supporting explanation because RoB table did not have a domain for attrition bias at all | 6 (0.06) |
| Total | 10292 (100) |

While there were too many examples of unclear explanations, we provide some examples of explanations categorized by us as unclear explanations in the Table 2.

**Table 2.**
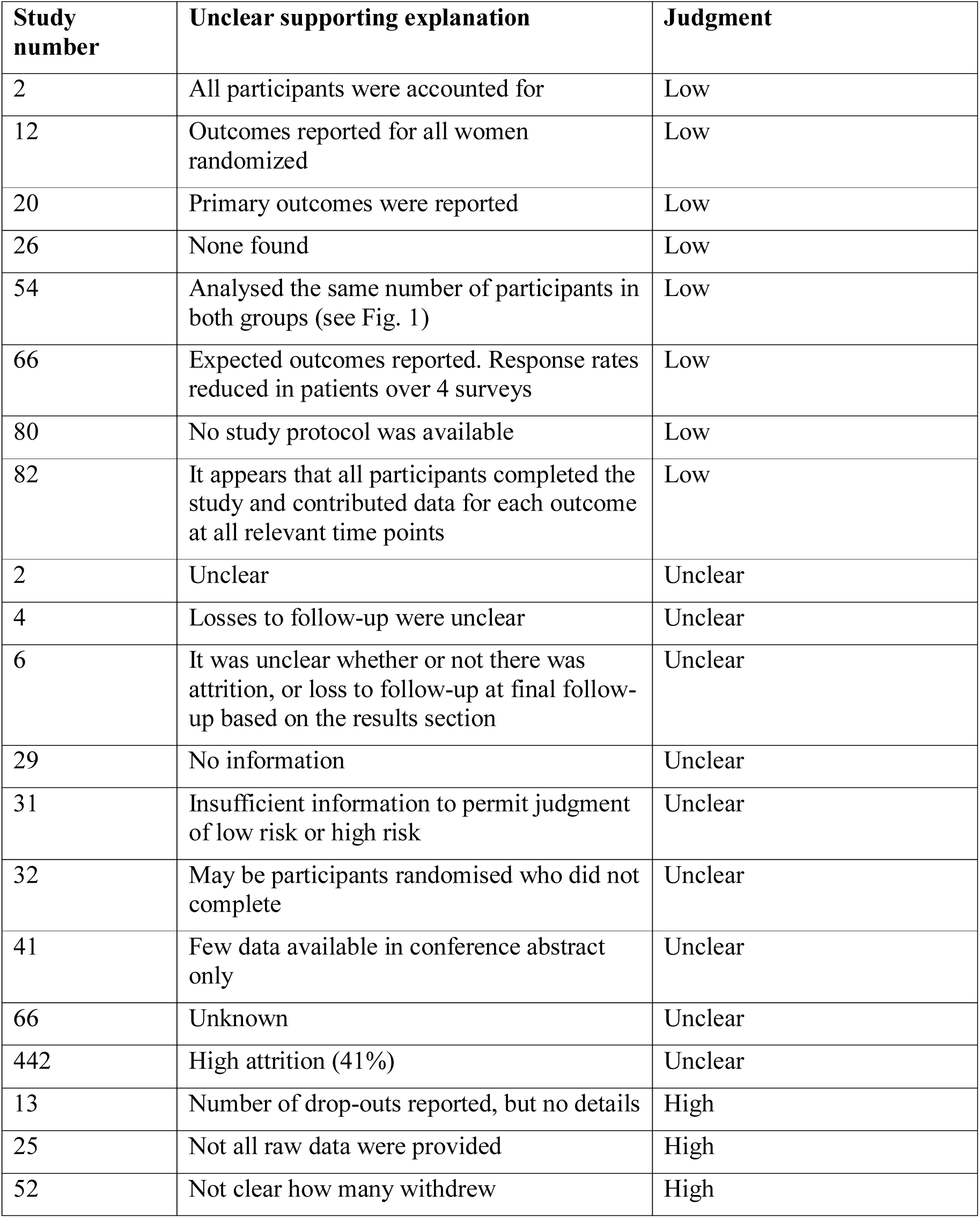
Examples of unclear supporting explanations

| **Study number** | **Unclear supporting explanation** | **Judgment** |
| --- | --- | --- |
| 2 | All participants were accounted for | Low |
| 12 | Outcomes reported for all women randomized | Low |
| 20 | Primary outcomes were reported | Low |
| 26 | None found | Low |
| 54 | Analysed the same number of participants in both groups (see Fig. 1) | Low |
| 66 | Expected outcomes reported. Response rates reduced in patients over 4 surveys | Low |
| 80 | No study protocol was available | Low |
| 82 | It appears that all participants completed the study and contributed data for each outcome at all relevant time points | Low |
| 2 | Unclear | Unclear |
| 4 | Losses to follow-up were unclear | Unclear |
| 6 | It was unclear whether or not there was attrition, or loss to follow-up at final follow-up based on the results section | Unclear |
| 29 | No information | Unclear |
| 31 | Insufficient information to permit judgment of low risk or high risk | Unclear |
| 32 | May be participants randomised who did not complete | Unclear |
| 41 | Few data available in conference abstract only | Unclear |
| 66 | Unknown | Unclear |
| 442 | High attrition (41%) | Unclear |
| 13 | Number of drop-outs reported, but no details | High |
| 25 | Not all raw data were provided | High |
| 52 | Not clear how many withdrew | High |

We categorized reported percent of attrition in the group with higher attrition into four categories: attrition under 10%, between 10 and 20%, between 21 and 30% and above 30%. Since some CSRs had multiple supporting explanations for a single judgment, we analyzed separately only CSRs where the only supporting explanation was about percent of attrition in the study groups. The purpose of this analysis was to see whether Cochrane authors use inconsistent judgments for various thresholds of attrition in this category of supporting explanations.

There were 264 CSRs that reported attrition that was under 10%. For 122 CSRs, this numerical category was the only explanation for the judgment. The majority of those CSRs judged difference in attrition between the groups that was under 10% as low risk of bias (101 CSRs, 82.8%), while 16 (13.1%) CSRs judged it as unclear risk of bias and 5 (4.1%) as high risk of bias.

Of 354 CSRs that reported attrition of 10-20%, 143 had this category as the only supporting explanation for the judgment. Of those 143 CSRs, 91 (63.6%) classified this as low, 28 (19.6%) as unclear and 24 (16.8%) as high risk of bias.

Among 215 CSRs that reported attrition of 21-30%, 60 had this category as the only supporting explanation for the judgment. Of those 60 CSRs, 34 (56.7%) judged this as low, 5 (8.3%) as unclear and 21 (35%) as high risk of bias.

There were 276 CSRs that reported attrition above 30%. For 70 this was the only category of supporting explanation, 18 (25.7%) classified this explanation as low, 9 (12.9%) as unclear and 43 (61.4%) as high risk of bias.

### Second category: difference in attrition between the groups

In the second category of supporting explanations about difference in attrition between the groups, 302/10292 (2.9%) explanations reported this category, and in all of them it was reported if the difference was above 10%.

### Third category: reporting of reasons for attrition

There were 2157/10292 (21%) supporting explanations related to reasons for attrition. The majority of these explanations referred to reasons for attrition that were reported in a trial, while the remaining supporting explanations indicated either that reasons for attrition were not reported in a trial, or that they were inadequately reported (Table 3).

**Table 3.** Number of supporting explanations in a category for reporting reasons for attrition

| **Category: reporting of reasons for attrition** | **N (%)** |
| --- | --- |
| Reasons reported | 1697 (16.5) |
| Reasons not reported | 370 (3.6) |
| Inadequately reported | 90 (0.9) |
| Total | 2157 (21) |

### Fourth category: supporting explanations about statistics

We found 1572/10292 (15.3%) supporting explanations related to statistics; Table 4 lists those that were mentioned in more than 5 CSRs in a way that they were described by the CSR authors themselves. Most of the explanations about statistics were referring to presence or absence of intention-to treat analysis (ITT), per protocol analysis (PP) or last observation carried forward (LOCF) (Table 4). Detailed analysis of risk of bias judgment categories was made only for the top five categories that reported only supporting explanation about statistics.

**Table 4.** Supporting explanations about statistics used that was related to attrition bias that were mentioned in more than 5 reviews

| **Statistics from the explanations** | **N (%)** |
| --- | --- |
| ITT | 825 (8) |
| No ITT | 238 (2.3) |
| PP | 88 (0.9) |
| ITT, LOCF | 86 (0.8) |
| LOCF | 66 (0.6) |
| ITT not reported | 47 (0.5) |
| ITT, PP | 34 (0.3) |
| mITT | 25 (0.2) |
| Completer analysis | 24 (0.2) |
| Sensitivity analysis | 15 (0.1) |
| BOCF | 12 (0.1) |
| ITT, BOCF | 8 (0.08) |
| Analysis not described | 6 (0.06) |

Acronyms: ITT = intention-to-treat analysis, LOCF = last observation carried forward, mITT = modified intention-to-treat analysis, PP = per protocol analysis

Of 825 CSRs that mentioned ITT in the explanation, there were 193 CSRs where this category of explanations was the only one. If a trial conducted ITT analysis, this was judged as low risk of bias in 140 (72.5%), unclear risk of bias in 21 (10.9%) and high risk of bias in 32 (16.6%) CSRs.

Among 238 CSRs where it was written that ITT was not used in a trial, in 35 this was the only category of supporting explanation. Of those 35 CSRs, 20 (57.1%) categorized this supporting explanation as low risk of bias, 9 (25.7%) as unclear and 6 (17.1%) as high risk of bias.

There were 81 CSRs that had PP analysis as a supporting explanation; 8 used it as the only explanation, with 7 (87.5%) that judged it low risk of bias and 1 (12.5%) that judged it as unclear risk of bias.

Of 66 CSRs that had LOCF as the supporting explanation for assessing risk of attrition bias, 25 CSRs had this as the only explanation; 13 (52%) judged this as a low risk of bias, 3 (12%) as unclear and 9 (36%) as high risk of bias.

There were 35 CSRs that indicated that it was unclear whether ITT analysis was used or not, because its usage was not described. None of those listed this item as the only supporting explanation for risk of attrition bias judgment.

### Inconsistencies in judgments in a single CSRs

We found only 34/729 (4.7%) CSRs that had inconsistencies in judging risk of attrition bias in the same CSR. This means that they gave different judgment for the same explanation. For example, “No incomplete outcome data” was judged as either low or unclear risk of bias in the CSR #210. In the CSR #255 explanation “No pre-publication protocol identified” was judged either as unclear or high. In the CSR #277 “No missing data” was judged as low or unclear. In the CSR #330 “No withdrawals mentioned” was judged as either low or unclear risk of attrition bias. There were 66/729 (9.1%) CSRs for which this analysis was not applicable because they included only one trial. All the other CSRs had consistent judgments for the given supporting explanations.

### Secondary analysis: Studies with multiple judgments of attrition bias for the same study

We found 27 CSRs that had multiple assessments of attrition bias for the same RCT. They had 2-7 multiple assessments separately, which we categorized in assessments related to aspects of attrition bias, time, objectivity and clinical outcomes.

Five CSRs had separate assessments of different aspects of attrition bias were assessments of drop-outs, participants analyzed in the group to which they were allocated and whether ITT analysis was performed. Seven CSRs had assessments related to time were multiple assessments for short-term or long-term outcomes, sometimes defined with specific time-frame (i.e. before or after 12 weeks or childhood outcomes), or end-of-intervention and end of follow-up. Five CSRs had separate assessments for subjective and objective outcomes. One of them specified what was a subjective and what an objective outcome was. Ten CSRs had separate assessments for different clinical outcomes (Table 5). The review authors did not analyze all these sub-domains for all studies included in those CSRs.

**Table 5.** Description of domains in Cochrane reviews that had multiple separate domains for assessing attrition bias for different outcomes

| **Study number** | **Names of separate domains for attrition bias in the Risk of Bias table** |
| --- | --- |
| 158, 197 | Short-term, long-term |
| 240 | End-of-intervention, end of follow-up |
| 250, 459, 533 | Subjective outcome measures, objective outcome measures |
| 285 | Clinical heart failure, subclinical heart failure (dichotomous and/or continuous), overall survival, tumor response, quality of life, adverse effects, adverse effects other than cardiac damage |
| 302 | Drop-out rate described and acceptable, participants analyzed in the group to which they were allocated |
| 312 | Mortality (all cause), hospital readmissions (all cause), hospital readmissions (due to adverse drug events), hospital emergency department contacts (all-cause), hospital emergency department contacts (due to adverse drug events), adverse drug events |
| 316 | Adverse events: hypothyroidism, development or worsening of Graves' ophthalmopathy, health-related quality of life, participants in euthyroid state, recurrence of hyperthyroidism, socioeconomic effects |
| 324 | 12 weeks or less, after 12 weeks |
| 340 | Primary outcomes, secondary outcomes |
| 346 | All outcomes: drop-outs, all outcomes: ITT analysis |
| 394 | Time to resolution of diabetic ketoacidosis, all-cause mortality, hypoglycemic episodes, morbidity, socioeconomic effects |
| 427 | Drop-outs reported, ITT analysis reported |
| 499 | Objective outcome (deaths), subjective outcome (quality of life) |
| 638, 795 | Drop-outs, ITT analysis |
| 641 | Pain, function |
| 722 | Short term follow-up (up to 3 months), longer term follow-up |
| 761 | Consumption outcome, selection outcome |
| 805 | Hemodynamic data, clinical outcomes |
| 867 | Survival, tumour response, toxicity, quality of life |
| 943 | Short-term outcomes, childhood outcomes |
| 946 | All outcomes, ITT analysis |
| 949 | Wound healed, wound area, time to healing |
| 951 | Pain, swelling, function, adverse effects |

### CSRs that did not have a domain for attrition bias in the RoB table

There were 12 Cochrane reviews that did not have a domain for attrition bias at all in the RoB table. They were not included in the main analysis, and hereby we report characteristics of their RoB tables. Five CSRs analyzed only 1 RoB domain, and this was ‘Allocation concealment in four cases (CSRs #341, #465, #672 and #904) and ‘Method for selecting cases to adjudicate?’ in one case (CSR #269). One CSR analyzed 3 RoB domains (Random sequence generation, Allocation concealment and Blinding as one domain for all outcomes), but not attrition bias (CSR #294). Three CSRs analyzed 4 RoB domains; one of them analyzed ‘Random sequence generation’, ‘Allocation concealment’, ‘Blinding of outcome assessment’, ‘Selective reporting’ (CSR #585) and two analyzed domains for ‘Random sequence generation’, ‘Allocation concealment’, ‘Blinding of participants and personnel (performance bias)’, ‘Size’ (CSRs #924, #936). Two CSRs analyzed five RoB domains (CSRs #174, #947) and one analyzed six RoB domains – but none of the domains were attrition bias (CSR #309).

### Risk of bias assessed with ‘yes’ or ‘no’ judgments

In 4/730 CSRs (0.5%) there was no standard judgment of risk of bias as high, unclear or low; instead RoB was judged as yes, unclear, no, or yes/no (CSRs #212, #292, #830 and #884). In one CSR risk of bias was graded as “low, unclear or high”, but in the supporting explanation also rated as A – Adequate, B – Unclear, C – Inadequate (CSR #244).

### Other inconsistencies that were encountered

Several CSRs had different name of the relevant domain. In the CSR #641 the domain was called “Intention-to-treat analysis performed?”, in the #419 “Losses to follow-up taken into account?” and in the CSR #873 “Complete follow-up?”.

### Explanations that should not be used for judging attrition bias

Finally, we decided to report examples of curious explanations for attrition bias judgments in Table 6. It appears to us that such explanations should not be used for explaining risk of attrition bias judgments.

**Table 6.** Examples of curious supporting explanations for attrition bias judgments that may not appear to be suitable for judging this risk of bias domain

| **Study number** | **Support for judgment** | **Judgment for risk of attrition bias** |
| --- | --- | --- |
| 82 | Chinese article - unable to ascertain | Unclear |
| 144 | This study was a feasibility study. Only 1 woman received the intervention. This study contributed no data to the review. | Unclear |
| 255 | No pre-published protocol identified | High or unclear |
| 256 | If we assume a person works for 40 hours per week, then for 28 participants the working hours will be 8960 hours for 8 weeks (4 weeks intervention and 4 weeks control period). However the study reported only 7,729 working hours based on accelerometer data | High |
| 376 | This is not clear from the paper. Author contacted, but when he moved jobs, the data files for this study were deleted | Unclear |
| 490 | 137 minus 28 equals 109, not 108 | Unclear |
| 492 | Exact time periods of 'before and after' accident data were unclear. Authors reported that they "should be 3 to 5 years". | Unclear |
| 494 | 1 - A reasonable account of how attrition was dealt with is given, but no specific reference to CONSORT | Low |
| 517 | Documented evidence that the CONSORT guidelines have been followed | Low |
| 606 | Data sparse largely narrative style | Unclear |
| 699 | Numbers do not always add up - query if N for outcomes are based on those who answered specific questions on follow-up? | High |
| 727 | Data of drop-outs was censored. | Low |
| 730 | Eleven patients were withdrawn before random assignment: 1 declined further participation, 8 were withdrawn by their physician, and 2 did not meet the entry criteria | Low |
| 744 | Publication is in German and our translation is incomplete. | Unclear |
| 835 | Differences in baseline characteristics of questionnaire responders vs non-responders (western ethnicity in 81% vs 54%, mean age 31 vs 28 years, median blood loss 1500 vs 1150 mL). Big difference in compliance to allocated treatment: 8 vs 34. The design of this trial carries a high risk for selecting the study population | High |
| 838 | Primary and secondary endpoints not specified directly but do address aims | Low |
| 849 | "The situations to consider eliminating the subject from data analysis did not arise" | Low |
| 850 | No Table 1 to clearly describe participant characteristics. | High |
| 854 | Duration of study not defined | High |
| 854 | Criteria for kidney disease not defined | Unclear |
| 873 | Denominators inconsistent in study | Unclear |

## Discussion

We found high inconsistency in the assessment of risk of bias related to incomplete outcome data, i.e. attrition bias in Cochrane systematic reviews. Cochrane authors do not have uniform approach to judging attrition bias. We did not observe clear numerical rules about the percent of attrition in trial groups or clear rules about statistics that was used or not used, that were consistently labeled as low, unclear or high risk of bias. One third of CSR authors had more than one category of explanations; some had up to four different categories. Inconsistencies were found even with the number of judgments, names of risk of bias domains and different judgments for the same explanations in the same CSR.

Cochrane Handbook indicates that “Missing outcome data, due to attrition (drop-out) during the study or exclusions from the analysis, raise the possibility that the observed effect estimate is biased.” The term ‘attrition bias’ is used for both exclusions and attrition [3]. Besides numerical indicators of attrition – absolute numbers and frequencies – that provide information about the magnitude of attrition, in the context of this domain of risk of bias different statistical methods for imputing missing data are often mentioned. For example, trial authors can use ITT analysis, or a ‘modified ITT analysis’. However, it has been reported that the term ‘ITT analysis’ does not always have a clear and consistent definition, and that it is not consistently used in trial reports [4]. The same was concluded for the modified ITT analysis and therefore it has been recommended by the Cochrane Handbook that the review authors should always ask information about who exactly was included in such analysis [3].

Simple imputations, such as last observation carried forward (LOCF) remain very popular despite warnings of statisticians against their use [5].

Judgments about different statistical methods varied in our analysis; we found very inconsistent judgments for different statistical methods. If we want to judge by the frequency of statistical comments in reviews where this was the only available explanation, we could not reach any conclusion, because the majority of authors judged presence of ITT analysis with low risk of bias, but also in the group that reported explicitly that there was no ITT analysis, this absence of ITT analysis was also predominantly judged with low risk of bias. Using *per protocol* analysis was mostly judged as low risk of bias, as well as LOCF analysis.

It has been published previously that attrition under 5% is not likely to introduce bias, while attrition rates above 20% raise concerns about the study validity [6]. While Cochrane handbook does not give clear guidance about the total attrition or attrition per group regarding specific numerical values, there is an example “17/110 missing from intervention group (9 due to ‘lack of efficacy’); 7/113 missing from control group (2 due to ‘lack of efficacy’)“ that is judged as high risk [3] in this example the first group has attrition of 15%. If a Cochrane author should follow this example, than attrition that is 15% or above per group should be labeled as high risk of bias.

In our study we found that numerical indicators for what represents attrition were widely inconsistent. When we categorized reported percent of attrition in the group with higher attrition and which threshold was predominantly judged in a certain way, attrition in a group that was under 10% was judged as low risk of bias in 83% of the cases, attrition 10-20% was judged as low risk of bias in 64% of cases, attrition 20-30% was judged as low risk of bias in 57% of cases. If we judge from the majority opinion of Cochrane authors, threshold of ‘above 30%’ is considered predominantly high risk of bias because 61% of judgments indicated so in CSRs where this was the only judgment so we could isolate the effect of this category for the overall judgment.

As for the risk of bias as a tool, it has been reported that it has low reliability between individual reviewers and across consensus assessments of reviewer pairs [7]. It has been argued that low reliability of the RoB assessment can have negative effects on decision making and quality of health care [8]. It has also been shown by da Costa et al. that standardized intensive training on RoB assessment may significantly improve the reliability of the Cochrane RoB tool [9]. However, our study points out that we would need first to have standardized instructions about what situations really represent risk of attrition bias. Having clear instructions, such as “attrition above 20% represents high risk of attrition bias” it would be much easier to achieve higher reliability of RoB assessment, even without formal training.

Instructions for assessing risk of attrition bias should include specific indications about all categories of assessment that should be appraised. It should be clearly specified which of those categories systematic review authors should assess, such as four that we used in this manuscript, including percent of attrition per group and difference between the groups, whether reasons for attrition were reported or not, and what is the appropriate statistics for dealing with attrition. If the authors do not have clear guidance about assessment of attrition bias, they can behave as we found – they can use one or more of those categories for their attrition RoB assessment as they personally see fit.

Some authors used multiple judgments for different follow-ups or different outcomes. This also introduces inconsistency in the attrition RoB assessment. Just as the option for authors to change the titles of attrition RoB domains in the RoB table in a Cochrane review.

Future studies on this topic should explore how to reduce inconsistency in assessment of attrition RoB, and they should attempt to reach consensus about what exactly should be assessed in this RoB domain.

In conclusion, we found very high inconsistency in methods of appraising risk of attrition bias in recent Cochrane reviews. Systematic review authors need clear guidance about different categories they should assess and judgments for those explanations. Clear instructions about appraising risk of attrition bias will improve reliability of the Cochrane risk of bias tool, help authors in making decisions about risk of bias and help in making reliable decisions in healthcare.

## Declarations

**Ethics approval and consent to participate:** Not applicable (secondary study of published manuscripts)

**Consent for publication:** Not applicable

**Availability of data and material:** Any additional information that were not presented in the manuscript are available on request from study authors

**Competing interests:** None

**Funding:** No external funding.

**Authors’ contributions:** Study design: LP, Data acquisition, analysis and interpretation: AB, RT, JASC, IN, JS, MV, IM, IV, TPP, LP. Writing of the first draft: LP, AB. Revising first draft for important intellectual content: AB, RT, JASC, IN, JS, MV, IM, IV, TPP, LP. Approval of the final version, and agreeing to be accountable for the work: AB, RT, JASC, IN, JS, MV, IM, IV, TPP, LP.

## Acknowledgements

None

